# Chronic stress facilitates behavioral engagement and alters lateral habenula activity during flexible decision making in a sex-dependent manner

**DOI:** 10.1101/2025.09.24.678366

**Authors:** Hayden R. Wright, Zachary D.G. Fisher, Ryan M. Schmid, Sara R. Westbrook, Riana A. Abeshima, Giuseppe Giannotti, Ryan J. McLaughlin

## Abstract

The ability to integrate feedback and flexibly adjust behavior under shifting environmental demands is required to optimize decision-making strategies. Clinical and preclinical data indicate that individuals with stress-related disorders and rodents exposed to chronic stress exhibit impaired behavioral flexibility. The lateral habenula (LHb) has emerged as a key brain region contributing to the effects of stress on cognitive performance. However, the extent to which the LHb is recruited to fine-tune decision-making strategies, as well as the impacts of chronic stress on LHb recruitment during task performance, remain largely unknown. To this end, we used a three-week model of chronic unpredictable stress (CUS) and performed *in vivo* fiber photometry to investigate Ca^2+^ transients in LHb neurons during an attentional set-shifting task in adult male and female Sprague Dawley rats (n=7-12/sex/group). We found that CUS exposure did not significantly impair behavioral flexibility. Rather, CUS-exposed rats made fewer omissions and exhibited shorter response latencies compared to controls, suggesting enhanced task engagement. We also observed sex differences in LHb Ca^2+^ activity. In control animals, we found that male rats showed stronger LHb signal prior to decision making, and greater activation following trial outcome than females. These differences were normalized by CUS, resulting in similar signaling patterns across sexes. Altogether, these findings reveal that chronic stress alters LHb activity in a sex-dependent manner without overtly impairing behavioral flexibility, thereby underscoring the importance of the LHb in decision making under stressful conditions.

**Highlights:**

- Chronic stress decreased response latency without impairing behavioral flexibility
- Male rats displayed greater inhibition of LHb activity prior to decision making
- Chronic stress abolished sex differences in LHb activity during decision making

## Introduction

The ability to incorporate feedback and adapt behavior to changing environmental demands is essential for effective decision-making. Importantly, dysfunction of cognitive flexibility is an often-overlooked feature of stress-related psychiatric disorders, including major depressive disorder (MMD). People with depression tend to be slower and less accurate in working memory tasks (Rose and Ebmeier, 2006; Harvey et al., 2004) due to decreased speed and attention, possibly resulting from reduced motivation or effort (Egeland et al., 2003). Flexible decision-making is also impaired in individuals with depression (Syder, 2013). Patients with depression struggle with cognitive flexibility during the Wisconsin Card Sorting Task, a widely used test in humans. After learning the initial rule, they often fail to apply it consistently, instead reverting to previously successful but now ineffective strategies, completing fewer categories than control groups within the allotted time (Grant et al., 2001). Stress directly contributes to deficits in flexible decision-making in humans (Hammen, 2005) and rodents (for review, see Hurtubise and Howland, 2017), likely through stress-induced alterations in mesolimbic and/or corticostriatal circuits that regulate various aspects of these complex decision-making processes. However, it remains unclear how the brain rapidly integrates changing contingencies, such as external cues (e.g., failing to find food) and internal states (e.g., increasing hunger), to modify ongoing behavior, and how stress impacts this process.

The LHb is an essential node in the circuitry underlying goal directed behavior, integrating environmental, internal, and rewarding or aversive feedback signals to influence reward-guided behavior via downstream dopamine, serotonin, and norepinephrine systems (Matusmoto and Hikosaka, 2007; Baker et al., 2017). The LHb is known to monitor ongoing behavioral states, influence decision making, and support the retrieval of spatial memories (Baker et al., 2015; Mathis et al., 2015). For example, pharmacological inactivation of the LHb impairs behavioral flexibility in rats trained to use auditory cues to choose between two arms in a maze, as indicated by an increased in perseverative errors committed when cues are switched in subsequent trials (Baker et al., 2017). Moreover, inactivation of the LHb abolishes reward preferences in a probabilistic choice task where rats can select a lever for either a small certain reward or a larger reward with decreasing odds of delivery over successive trial blocks (Stopper and Floresco, 2014). Together, these findings suggest that the LHb uses both proactive (cue-based) and retroactive (reward outcome-based) information to guide future choices and plays a critical role in evaluating reward magnitude when outcomes are uncertain, implicating it as a key regulator of reward-guided decision making.

Importantly, LHb dysfunction has been linked to behavioral flexibility deficits and maladaptive decision making observed in humans with stress-related disorders such MDD and in rodents exposed to chronic stress. Humans with MDD and rodents subjected to a chronic unpredictable stress (CUS) regimen that models core features of MDD show LHb hyperactivity (Morris et al., 1999; Berger et al., 2018; Cerniauskas et al., 2019, Zhang et al., 2023).Clinical studies have shown that normalizing LHb firing via deep brain stimulation can significantly alleviate symptoms in patients with treatment-resistant depression (Sartorius et al., 2010; Wang et al., 2021) and lessens depressive-like behaviors in rodents (Li et al., 2011). Although acute stress impacts LHb reward signaling (Shabel et al., 2019) and basic reward-guided decision making in a T-maze task (Nuno-Perez et al., 2021), the effects of chronic stress on LHb dysfunction in relation to behavioral flexibility have yet to be examined. Many studies have documented consistent effects of chronic stress on different aspects of flexible decision-making in males (Hurtubise and Howland, 2017). However, no research has investigated how chronic stress influences operant-based attentional set-shifting performance, or how this relates to stress-induced LHb dysfunction. The present study addresses this gap by examining how a history of CUS exposure alters LHb activity during flexible decision making, providing new insight into the circuit-level mechanisms underlying stress-induced deficits in behavioral flexibility.

### Experimental Procedures

#### Subjects

Male (n=38) and female (n=38) Sprague-Dawley rats were purchased from Envigo and arrived at postnatal day 65. Animals were pair-housed in cages approximately 35cm x 45cm x 20cm. Cages were housed in a humidity and temperature-controlled vivarium maintained on a 12-hour reverse light cycle (7:00 lights off, 19:00 lights on) with ad libitum food and water access. All testing occurred under red light during the active cycle (lights off). Animal care adhered to the National Research Council’s Guide for the Care and Use of Laboratory Animals and Guidelines for the Care and Use of Mammals in Neuroscience and Behavioral Research, with all procedures approved by the Washington State University Institutional Animal Care and Use Committee.

#### Intracranial surgery

Rats were anesthetized with 2-5% isoflurane in 1L/min O_2_ and mounted into a stereotaxic frame (**Fig. 1**). All animals were unilaterally injected with 300nL of AAV1.Syn.GCaMP6s.WPRE.SV40 virus (1×10^13^ vg/mL, Addgene, Watertown, MA) into the LHb (AP -3.6mm; ML +/-0.7mm, DV -5.0mm) before implantation of the optic fiber cannula (200 mm core, 0.37 NA, Neurophotometrics, San Diego, CA, USA) 0.2mm dorsal from the injection site. The injector was allowed to rest 2 min after lowering to the injection site before the infusion was administered over 3 min. The injector was then left in place for 10 min to allow for diffusion into the tissue and minimize viral spread up the injection track. Following infusion, four stainless-steel jeweler’s screws were implanted surrounding the injection site. The fiber optic cannula was placed and secured to the skull with ultraviolet light-curable dental acrylic (FusionFlo, Prevest Denpro, Jammu, India) using the jewelers screws as anchors. Immediately following surgery and each day for the following three days, rats were weighed and received antibiotics (enrofloxacin, 5mg/kg) in their drinking water, and oral analgesics (Meloxicam, 1mg/kg). All rats were individually housed and given 10 days of recovery before resuming group housing.

**Figure 1:**
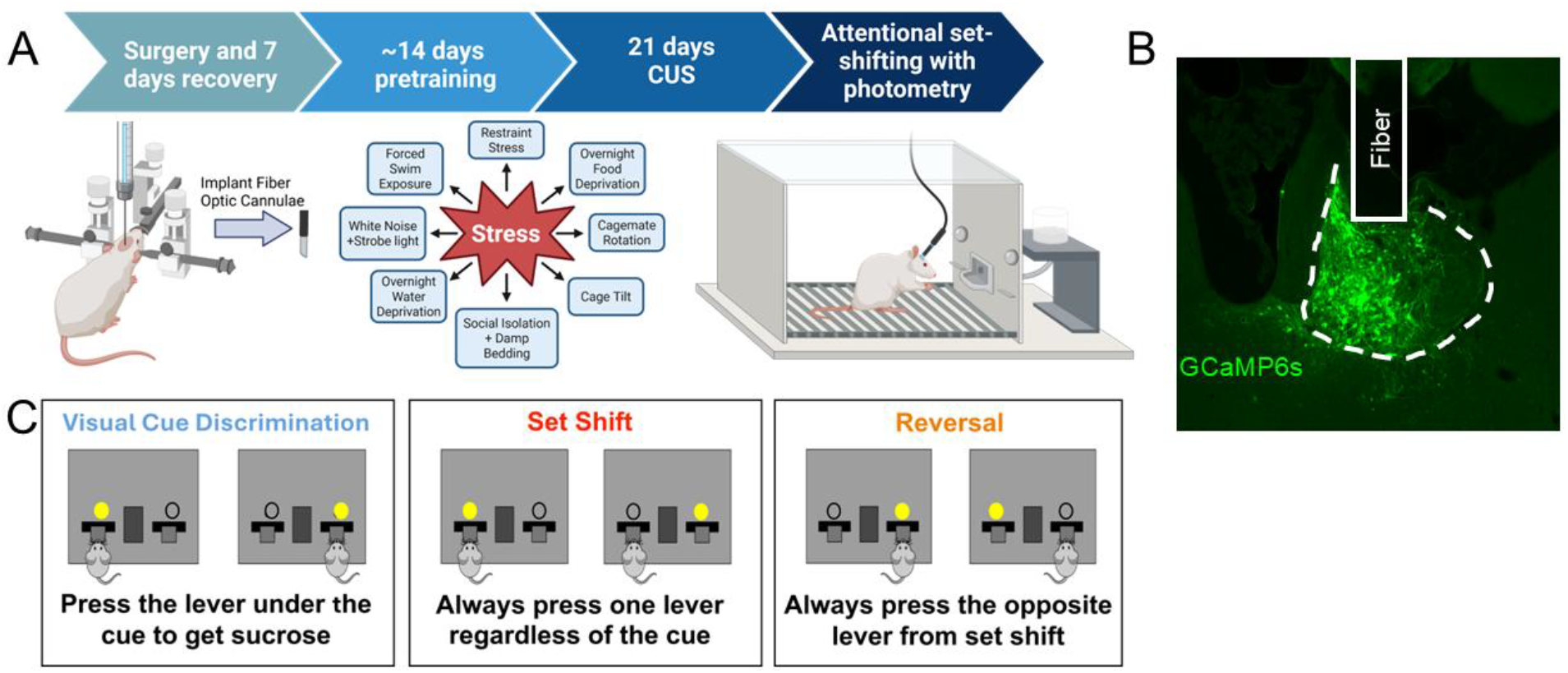
Experimental Details. **(A)** Experimental timeline and strategy. **(B)** Representative image of GCaMP6s expression and fiber optic cannulae placement within the LHb (dotted line). **(C)** Visual explanation of behavioral flexibility task. Created in BioRender. Winuthayanon, J. (2025) https://BioRender.com/ddoohgz Used with permission.

#### Chronic unpredictable stress (CUS)

A modified version of the CUS paradigm was used, as described previously (McLaughlin et al., 2013; Berger et al., 2018). Briefly, the CUS paradigm consisted of three weeks of unpredictable exposure to two or three daily mild stressors from the following list: immobilization using Broome restrainers (30 min), socially crowded open field exposure with white noise/strobe light (1 hr), cage rotation to alter dominance hierarchies (3 hr), social isolation with wet bedding (18 hr), forced swim exposure (5 min), overnight food or water deprivation (18 hr), and 45° cage tilt (45 min). Rats were not deprived of food or water overnight while undergoing the testing phase of the attentional set-shifting task.

#### Attentional set-shifting task

Five days before pretraining, baseline weights were recorded, and rats underwent food restriction until they reached 90% of their free-feeding baseline, which was then carefully maintained throughout pretraining. This food restriction procedure was repeated near the end of the CUS period, at which point a new baseline measure was obtained. Attentional set-shifting was conducted in 10” x 12” x 12” (L x W x H) Coulbourn Habitest chambers (Coulbourn Instruments, Holliston, MA) using Graphic State 4 software. A full description of the operant training process is described in (Weimar et al. 2020) and (Freels et al. 2024), which was modified from (Floresco, Block, and Tse, 2008; Brady and Floresco, 2015). Once all animals housed in a cage completed pretraining, they began a period of either three weeks of CUS or no stress. During this period, animals received a weekly pretraining reminder session on the retractable lever task without food restriction. CUS was maintained throughout the attentional set-shifting procedure, though on test days rats were not exposed to stressors until after their recording session.

Next, rats advanced to the visual cue discrimination, set shift, and reversal testing phases. In each task, a cue light was illuminated for three seconds before lever presentation, at which point the rat had 10 seconds to respond, otherwise an omission was scored and the levers retracted. The reward was delivered or withheld immediately following the lever press response. Criterion for completion of each task was 10 correct consecutive responses during a session. In the visual cue discrimination task, rats were required to press the lever directly below an illuminated cue light to receive a 45 mg sucrose pellet reinforcer. During the set-shifting task, an extradimensional shift was introduced, such that rats had to disregard the previous visual cue-based rule and instead press the lever opposite of their side preference, which was determined during pretraining. For the reversal task, rats were required to make an intradimensional shift by learning to press the lever opposite the lever that was reinforced during the set-shift task to receive sucrose reinforcement. Daily sessions ended when criterion was met or after 200 trials. Assessment of behavioral flexibility was based on the total number of trials rats needed to reach criterion during set-shift and reversal tasks. Error types were analzyed for set-shifting and reversal tasks by dividing sessions into 16 trial blocks as described (Brady and Floresco 2015). In the set-shifting task, errors made following the previous strategy (visual cue) were scored as perseverative until fewer than six errors were made in a block, at which point errors were classified as regressive. Never-reinforced errors were classified when a rat made a response that was not reinforced in either the visual-cue discrimination or set-shifting task. In the reversal task, errors were classified as perseverative until fewer than 10 errors were made in a block, with subsequent errors then scored as regressive. Whether the errors were made toward or away from the cue light were also recorded.

#### Fiber photometry recordings

Ca^2+^ dependent fluorescence (470nm) and an interleaved isosbestic control wavelength (415nm) were recorded during each test day at 100Hz using a branched fiber optic patch cord (Doric, Quebec City, Canada), a FP3002 acquisition system (Neurophotometrics, San Diego, CA, USA), using BonsaiRX software (NeuroGEARS, London, United Kingdom) with simultaneous timestamping at the time of lever press via a TTL pulse. Rats were acclimated to the recording procedure for five min daily for three days prior to the first test day. Rats were under continuous observation during photometry data collection.

Analysis of photometry data was performed as in Martianova, Aronson, and Proulx (2019) using custom Python scripts. Decision-making events were identified as correct or incorrect with timestamps from TTL pulses from the operant chamber system. Each event was defined as the section of recording five seconds before the response and eight seconds afterwards. This window guarantees that no overlap of trials is possible. The pre-response phase was defined as three seconds before the lever press to capture LHb activity during cue light presentation up to the point in time the response was made. The post-response phase was defined as the four seconds after the lever was pressed to capture the LHb signal during the anticipation of and reaction to either reinforcer delivery or lack thereof. Signals around correct and incorrect trials were concatenated into lists for each task for each rat. Area under the curve (AUC) was calculated using the trapezoidal rule for pre- and post-response periods on every trial in each task, which was then used to obtain an average for each subject.

#### Brain preparation and histology

Rats were anesthetized with an intraperitoneal injection of ketamine (91mg/kg) and xylazine (9.1mg/kig), transcardially perfused with ice-cold phosphate-buffered saline (PBS) for two minutes, then 4% paraformaldehyde for 6-10 minutes before brain extraction. Brains were kept in 4% PFA overnight before being transferred to PBS containing 30% sucrose and 0.1M sodium azide for 72 hr. Brains were then flash frozen and stored at -80C. Brain slices (30 µm) containing the LHb were collected using a cryostat (Leica Biosystems, Buffalo Grove, Illinois, USA), mounted on superfrost plus slides (Thermo Fisher Scientific, Waltham, Massachusetts, USA), coverslipped (Electron Microscopy Sciences, Hatfield, Pennsylvania, USA), then imaged on an epifluorescence microscope (Leica Biosystems, Buffalo Grove, Illinois, USA). Rats with viral expression or implant placements outside the anatomical boundaries of the LHb were excluded from fiber photometry analyses but still included in behavioral analyses (Behavior: M CUS= 20, M CTRL=17, F CUS =15, F CTRL =20; Photometry: M CUS=12, M CTRL=9, F CUS=10, F CTRL=7).

### Statistical analysis

Statistical analysis was conducted using SAS software (version 9.4, SAS Institute Inc., Cary, NC, USA). For behavioral measures, two-way analysis of variance (ANOVA) was used with sex and stress as between subjects factors. Analysis of fiber photometry data was conducted using three-way ANOVAs, with sex and stress as between-subjects factors and response (i.e., correct vs. incorrect) as a within-subjects factor. All significant interactions were followed up with simple main effects analyses. Significant 3-way interactions were followed up with 2-way ANOVAs conducted separate by sex. In rare cases, fiber photometry AUC values greater than three standard deviations from the group mean were winsorized to three standard deviations to minimize the influence of extreme outliers without discarding useful data (n=10 of 456 included values.

## Results

### Chronic stress facilitates behavioral engagement during flexible decision making

Male and female rats were microinjected with an AAV containing a construct to express the fluorescent calcium indicator GCaMP6s into the LHb before implantation of a fiber optic cannula (**Fig 1A** and **1B**). After recovery and completion of pre-training, rats were exposed to three weeks of CUS, which we and others have used to elicit robust elevation of corticosterone, adrenal hypertrophy, and a depressive-like behavioral phenotype (Berger et al., 2018; McLaughlin et al., 2013; Hill et al., 2005; Hill et al., 2008; Reich, Taylor, and McCarthy., 2009). To assess behavioral flexibility, we used an operant based attentional set-shift-reversal task (**Fig. 1C**), and Ca^2+^-dependent fluorescence from the LHb was measured during the visual cue discrimination, set-shift, and reversal phases of the behavioral flexibility task.

No significant effects of CUS or sex were observed in the number of trials needed to complete the tasks during either the shift (**Fig. 2A**) or reversal (**Fig. 2E**) tests. Incorrect trials were classified by type (Brady and Floresco, 2015) and quantified for shift and reversal tasks. We found a main effect of sex on never-reinforced type errors during the shift task (F_1,67_= 4.73, p=.033), where males committed fewer errors of this type than females, irrespective of stress condition (**Fig. 2D**). No differences in types of errors were observed during reversal testing (**Fig. 2H**). While CUS did not impact behavioral flexibility performance, CUS-exposed rats unexpectedly committed significantly fewer omissions compared to control rats during visual cue discrimination (F_1,68_= 6.1, p=.0160, **Fig. S1B**), and shift (F_1,68_= 7.83, p=.0067, **Fig. 2C**), but not during reversal (**Fig. 2G**). CUS-exposed rats also exhibited a decreased latency to press after cue presentation during visual cue discrimination (F_1,68_= 7.94, p=.0063, **Fig. S1C**), shift (F_1,67_= 13.41, p=.0005, **Fig. 2B**), and reversal (F_1,68_= 10.63, p=.0017, **Fig. 2F**) compared to control rats. There were no group differences during acquisition of visual cue discrimination (**Fig. S1A**), or accuracy across shift (**Fig. S1D**) or reversal (**Fig. S1E**). Collectively, these data indicate that CUS increases task engagement without impairing learning or behavioral flexibility.

**Figure 2:**
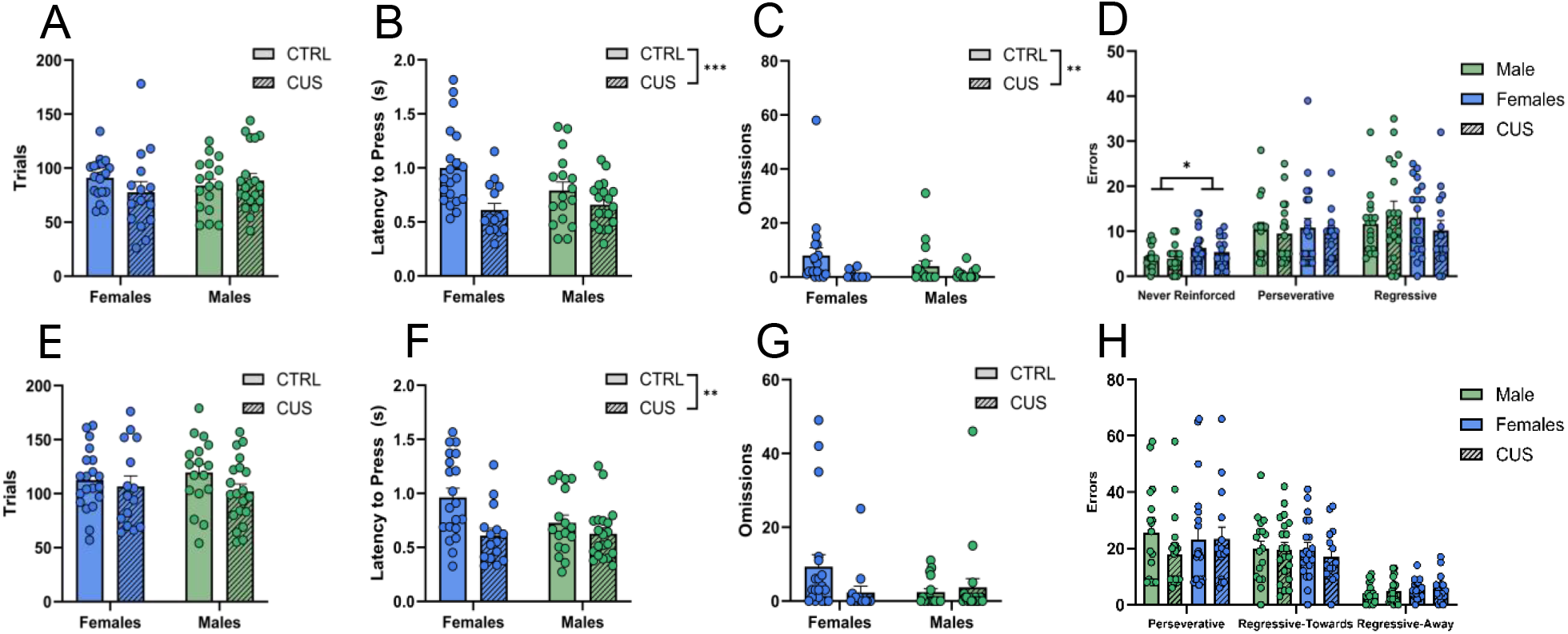
Chronic stress facilitates behavioral engagement without altering behavioral flexibility performance. **(A)** Trials to criterion in shift and **(E)** reversal. **(B)** Latency to press in shift and **(F)** reversal. **(C)** Number of omissions during shift and **(G)** reversal. **(D)** Number of error subtypes in shift: never-reinforced, perseverative, and regressive. **(H)** Number of error subtypes in reversal: perseverative, regressive-toward cue light, and regressive-away from cue light. All statistical tests are two-way ANOVAs with stress and sex as a factor. Simple main effect of stress group is denoted in the top right legend of each graph, where applicable. Main effects of sex were denoted in the figure, where applicable. *p< 0.05, **p<0.01, ***p<0.001

### LHb Ca^2+^ dynamics during flexible decision making are sex dependent and differentially altered by chronic stress

To assess differences in LHb Ca^2+^ dynamics between groups during correct and incorrect trials, AUC values were calculated before and after the lever press response. The pre-response phase was defined as three seconds before the lever press, which captured LHb activity during cue light presentation up to the point in time the decision was expressed. The post-response phase was defined as the four seconds after the lever was pressed, which captured LHb signal during the anticipation of, and reaction to, either reinforcer delivery or lack thereof.

During the set-shift component of the task, there were significant interactions of stress and sex in the pre-response period (F_1,34_=10.65, p=.0025), as well as the post-response period (F_1,34_=5.08, p=.031) (**Fig. 3A, 3B**). These interactions were examined using simple main effects testing, which indicated a main effect of sex within the control animals for both the pre-response (F_1,34_=12.29, p=.0013) and post-response (F_1,34_=10.35, p=0.0028) periods. During the pre-response period, control females exhibited a smaller decrease in AUC (**Fig. 3C**) than their male counterparts (**Fig. 3E**). In the post-response period, females showed a further decrease in AUC (**Fig. 3D**) whereas males showed a sharp increase (**Fig. 3F**). Notably, this sex difference was not present in CUS-exposed rats, whose AUC values were affected in opposite directions relative to controls, such that the CUS males and CUS females did not statistically differ. Additional simple main effect analyses revealed that, in males, CUS exposure attenuated the pre-response reduction in AUC compared to controls (F_1,34_=8.35, p=.0066). We also observed a main effect of trial outcome in the post-response phase of shift (F_1,34_=12.53, p=.0012), where all groups exhibit an increased AUC during incorrect trials compared to correct trials.

**Figure 3:**
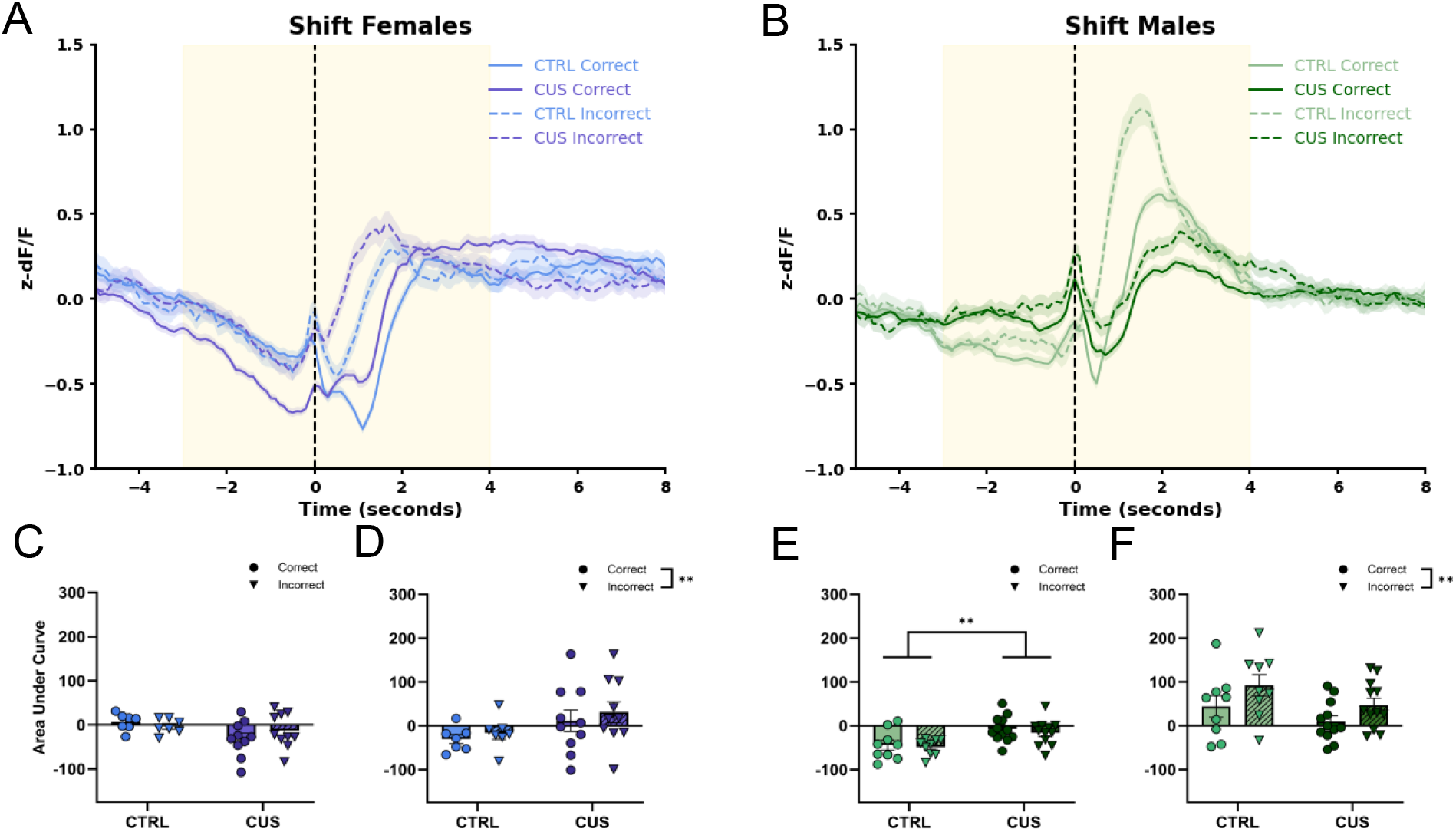
LHb Ca^2+^ dynamics during set-shift are sex dependent and differentially altered by chronic stress. Averaged group traces of shift-correct trials for **(A)** females and **(B)** males. Yellow shading indicates the data frames from which area under the curve (AUC) was calculated. AUC **(C)** pre- and **(D)** post-response for correct and incorrect trials during set-shift in females. AUC **(E)** pre- and **(F)** post-response for correct and incorrect trials during set-shift in males. Statistics are 3-way ANOVAs with sex, stress, and response type as factors. Main effects of response type are indicated in the legend in the top right of each AUC graph. Main effects of sex are denoted in the graph area. Main effects of stress are common and not indicated in the graph, only the body of the results. *p<0.05, **p<0.01, ***p<0.001

The reversal test was conducted the day after criterion is met on the shift task. Consistent with the set-shift data, during the reversal test there was a significant interaction of stress and sex on LHb activity during the pre-response phase (F_1,34_=9.09, p=.0048), as well as the post-response phase (F_1,34_=4.60, p=.039) (**Fig. 4A, 4B**). As in the prior test, female control rats exhibited more LHb activity (**Fig. 4C**) than control males during the pre-response period (**Fig. 4E**) (F_1,34_=11.97, p=.0015), and significantly less LHb activity during the post-response phase (**Fig. 4D**) (F_1,34_=8.02, p=.0077) than control males (**Fig. 4F**). CUS again led to a similar pattern of LHb activity, whereby activity was shifted to closely resemble the activity profile in the control group of the opposite sex. In the pre-response phase, there was a simple main effect of stress, with control females displaying significantly greater AUC than CUS females (**Fig. 4C**) (F_1,34_=6.83, p=.013). In addition, within both phases, there was a main effect of trial outcome. In the pre response phase, there was less Ca^2+^ activity recorded prior to selecting the incorrect lever (F_1,34_=4.54, p=.040), and in the post-response phase, there was more Ca^2+^ activity in response to an incorrect trial (F_1,34_=30.06, p<.0001). Effects similar to those seen in shift and reversal were also evident during visual cue discrimination (**Fig. S2**)

**Figure 4:**
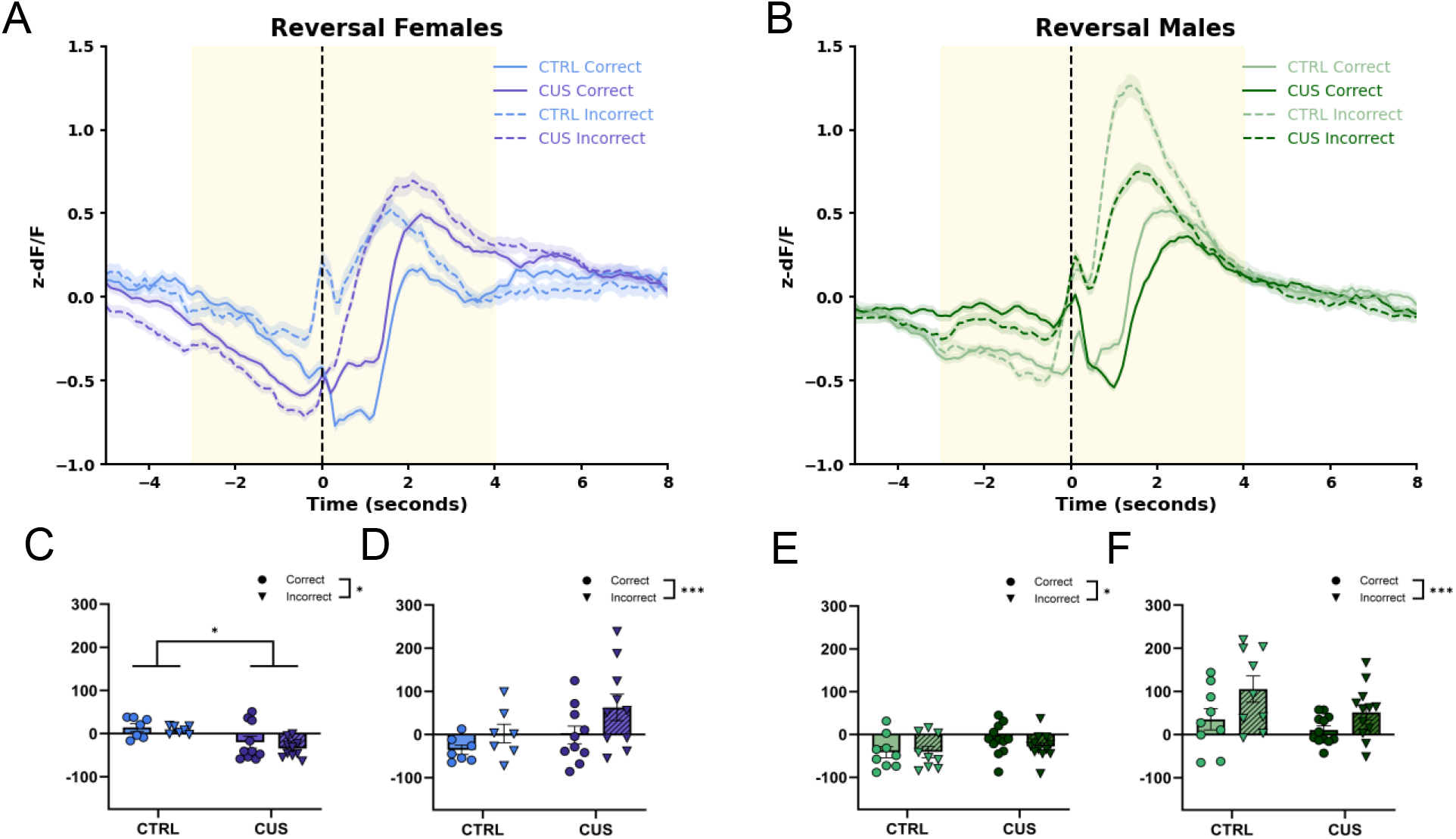
LHb Ca^2+^ dynamics during reversal learning are sex dependent and differentially altered by chronic stress. Averaged group traces of reversal-correct trials for **(A)** females and **(B)** males. Yellow shading indicates the data frames from which area under the curve (AUC) was calculated. AUC **(C)** pre- and **(D)** post-response for correct and incorrect trials during reversal in females. AUC **(E)** pre- and **(F)** post-response for correct and incorrect trials during reversal in males. Statistics are 3-way ANOVAs with sex, stress, and response type as factors. Main effects of response type are indicated in the legend in the top right of each AUC graph. Main effects of sex are denoted in the graph area. Main effects of stress are common and not indicated in the graph, only the body of the results. *p<0.05, **p<0.01,***p<0.001

## Discussion

Here, we investigated the effects of CUS on an operant-based attentional set-shifting task that models flexible decision making in male and female rats. Surprisingly, our data reveal that our CUS regimen did not impair set-shifting or reversal performance. Conversely, CUS-exposed rats made fewer omissions and exhibited shorter response latencies, consistent with enhanced task engagement (**Fig. 2**). In control rats, we observed sex differences in LHb activity, such that males showed larger pre-response inhibition and larger post-response activation than females. These differences were abolished following CUS, yielding more similar LHb signaling across sexes (**Figs. 3-4**). Together, these corresponding behavioral and physiological changes raise the possibility that stress-induced remodeling of LHb activity favors rapid, exploitative responding that preserves operant performance in subjects exposed to chronic stress. This, in turn, could help to buffer the deleterious effects of CUS on flexible decision-making.

To our knowledge, we are the first to assess the effects of CUS on behavioral flexibility in an operant context. The negative impacts of chronic stress generally, and CUS specifically, on both set-shifting and reversal learning are well documented (Bondi et al., 2008, 2010; Hill et al., 2005). However, in the present study we did not observe an effect of our CUS regimen on set-shift or reversal learning. Among the factors that may explain this incongruency, the type and specific features of the stress paradigm used often play a key role. It is important to note that other studies employed two-week CUS models (Bondi et al., 2008, 2010), or one and three weeks of chronic restraint stress (Nikiforuk 2012a, 2012b), which differs methodologically from the three-week CUS regimen used in our study. Another factor that might explain this behavioral incongruency is the task used to assess behavioral flexibility. Previous studies showing chronic stress-induced deficits in flexible decision-making used tasks that required using olfactory/tactile cues (Birrell and Brown, 2000) or spatial maze navigation (Baker et al., 2017), whereas the present study employs lever pressing guided by visual cues and egocentric spatial awareness. The strength of the operant approach is that it assesses behavioral flexibility independently of factors that can introduce confounds, such as innate olfactory preferences, repeated handling, or inadvertent sensory cues during manually administered trials (Floresco, Block, and Tse, 2008)). Nevertheless, it remains possible that these ethologically relevant cues or spatial navigation demands are precisely what recruit the LHb into the active ensemble guiding decision-making.

Given that CUS did not change the total number of trials required to complete the task, and experimenters observed less exploration of the operant chamber by CUS-exposed rats, we theorized that stress did not impair learning but instead enhanced task engagement. Consistent with this, CUS-exposed rats pressed the lever more quickly and made fewer omissions than control rats across all task phases. Our observations align with reports that rats subjected to five weeks of CUS collected rewards more rapidly than controls, driven by increased locomotion and reduced off-task behavior, while other aspects of decision-making remained unaffected (Matisz, Badenhorst, and Gruber, 2021). CUS-exposed rats also made fewer anticipatory licks in this task (Matisz, Badenhorst, and Gruber, 2021). Together, these findings suggest that enhanced task engagement and more efficient reward collection in an operant context occur independently of stress-induced negative biases in outcome prediction or hedonic valuation, which are consistently reported in stressed rodents (Shabel et al., 2019; Cerniauskas et al., 2019). Altogether, CUS may induce changes that optimize resource gathering over exploration in a non-threatening context. However, future studies are needed to thoroughly investigate and characterize the effect of stress on “exploitation vs. exploration” resource gathering strategies.

We used *in vivo* fiber photometry to monitor dynamic Ca^2+^-dependent fluorescence as a proxy for bulk LHb activity in male and female rats, that were either chronically stressed or not. Although Ca^2+^ signals do not always map directly onto spiking (Legaria et al., 2022), subthreshold fluctuations are known to support signal processing, plasticity, and intracellular signaling. Consistent with prior work, we observed increased LHb signals following incorrect trials compared to correct trials, reflecting the well-established activation of the LHb by aversive stimuli, such as omission of an expected reward (Matsumoto and Hikosaka, 2007; Shabel et al., 2019; Nuno-Perez et al., 2021). Notably, to our knowledge, this is the first demonstration of such responses in female rats.

While the magnitude of LHb responses to reward omission on incorrect trials aligned with our predictions, we did not anticipate that CUS would differentially affect male versus female LHb activity. Baseline LHb activity was modulated by CUS, such that female CUS rats resembled control males, and male CUS rats resembled control females. Previous studies report heightened LHb responses to reward omission following stress (Shabel et al., 2019; Nuno-Perez et al., 2021); however, during the shift component of the task, CUS-exposed males showed a blunted response relative to controls. This may reflect a limitation of fiber photometry, which measures fluorescence relative to session-specific baselines: a broad CUS-induced increase in LHb activity could obscure dynamic changes in task-responsive neurons.

The sex differences in LHb activity may stem from structural differences in LHb inputs. Human and rodent studies report sex-dependent variation in the size and number of habenular inputs (Hitti et al., 2022; Liu et al., 2022). Such differences could lead to different patterns of LHb activity and allow males and females to achieve similar behavioral outcomes via distinct mechanisms of downstream monoaminergic activation. Supporting this idea, we observed sex-specific error patterns during the shift phase, despite equivalent trials to criterion. Recent work shows reduced LHb-mediated inhibition of dopaminergic firing in females and less variable LHb firing patterns (Bell, Waldron, and Brown, 2023), and cell-type-specific encoding of reward and aversion is critical for feedback-guided decision-making (Mondoloni, Mameli, and Congiu, 2021; Shabel et al., 2019). Thus, sex differences in LHb inputs could produce different patterns of LHb activity that engage distinct monoaminergic pathways. More specifically, male LHb signaling might provide stronger inhibition of dopamine neurons after errors, whereas female LHb signaling might produce less inhibition and more sustained dopamine firing, which would collectively support our observation of similar behavioral outcomes occurring via distinct circuit mechanisms.

However, before drawing mechanistic conclusions, it is important to note some limitations of our experimental design that temper interpretation of the observed relationship between CUS, LHb signaling, and task engagement. As noted above, these data are correlational and reflect bulk LHb activity, so causality (and the precise cell-type or projection-specific mechanisms) remains to be established. Opposing changes in distinct LHb cell populations or projections could be responsible for different aspects of task performance that were obscured in the bulk signal. Additionally, it remains possible that CUS-driven shifts in tonic LHb activity could have altered AUC measures, or that effects of CUS on task performance are more driven by changes in arousal, locomotion, or outcome sensitivity. In future studies, causal manipulations (e.g., projection- or cell-type-specific perturbations) will be required to fully determine whether LHb changes are sufficient and necessary to drive the observed increases in task engagement.

Altogether, these findings indicate that chronic stress remodels LHb signaling in a sex-convergent manner that is associated with increased task engagement, reframing the LHb as a circuit node that may support adaptive behavioral strategies under conditions of chronic stress. By linking chronic stress, sex-specific LHb activity, and motivated responding, this work shifts the focus from viewing stress as solely detrimental toward recognizing circuit-level adaptations that could be targeted to promote resilience. More broadly, identifying LHb circuit signatures that distinguish adaptive from maladaptive stress responses could inform interventions to preserve cognitive function in stress-related neuropsychiatric disorders.

## Acknowledgements

The authors would like to thank Leisa Uelese and Ryan Faddis for their vital assistance in running behavioral tests and providing exceptional animal care.

## Author Contributions

RJM conceived the idea, designed the study, and edited the manuscript. HRW performed surgeries, conducted chronic stress exposure and behavioral testing, analyzed and interpreted the data, and wrote the manuscript. ZDGF performed daily colony maintenance and animal husbandry, performed surgeries, conducted chronic stress exposure, conducted behavioral testing, analyzed data, and edited the manuscript. RMS performed surgeries, conducted chronic stress exposure, conducted behavioral testing, and data analysis. SRW assisted with colony maintenance and animal husbandry, conducted chronic stress exposure, and data analysis. RA assisted with colony maintenance, animal husbandry, and behavioral testing. GG provided data analysis pipelines for fiber photometry analyses, helped to analyze and interpret the data, and edited the manuscript.

## Supplemental Figures

**Supplemental Figure S1:**
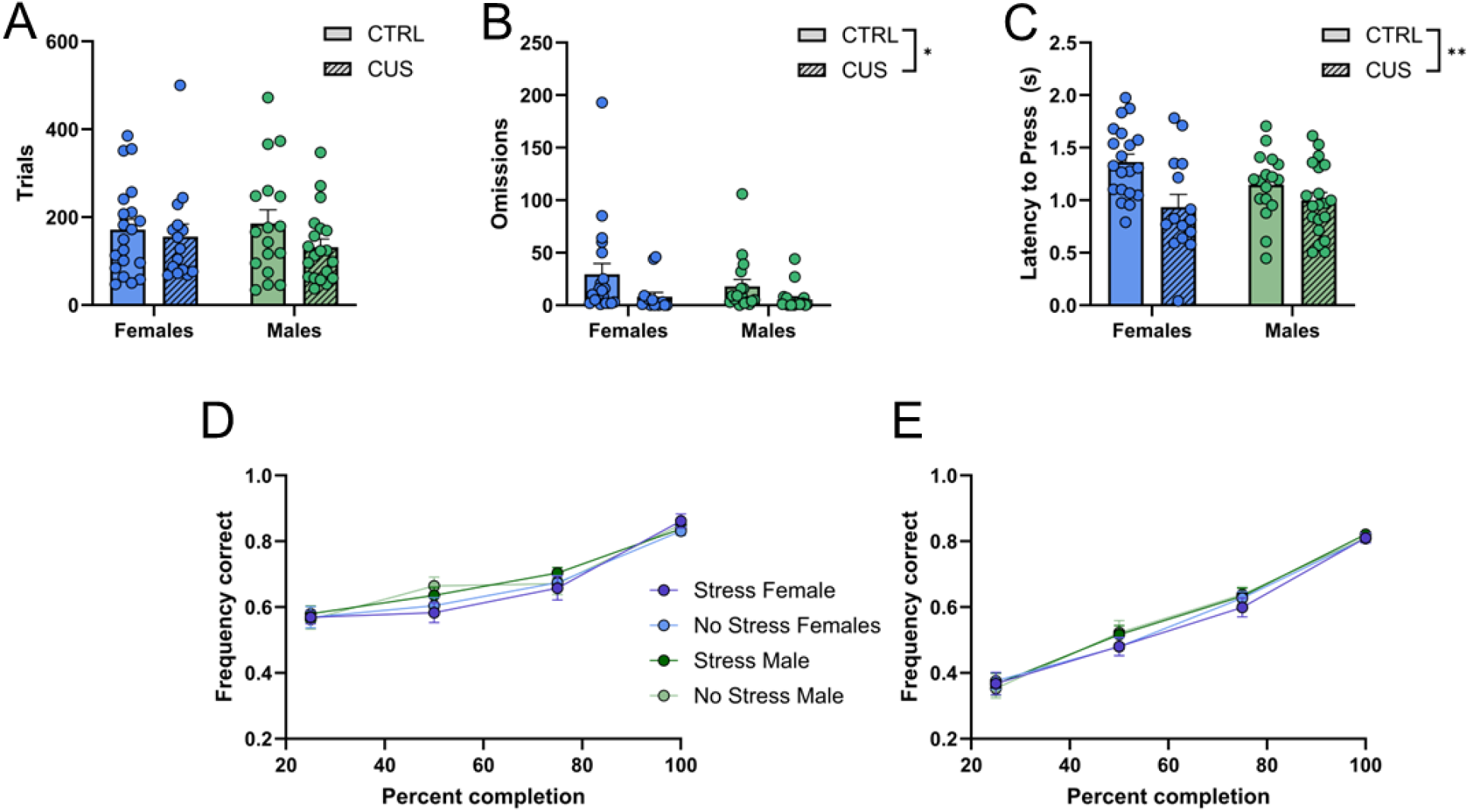
Chronic stress facilitates behavioral engagement without improving visual cue discrimination performance or learning rate during behavioral flexibility. **(A)** Trials to criterion, **(B)** number of omissions, and **(C)** latency to press during visual cue discrimination. Accuracy during **(D)** shift and **(E)** reversal. All statistical tests are two-way ANOVAs with stress and sex as a factor. Simple main effect of stress denoted in top right legend of each graph, where applicable. *p< 0.05, **p<0.01.

**Supplemental Figure S2:**
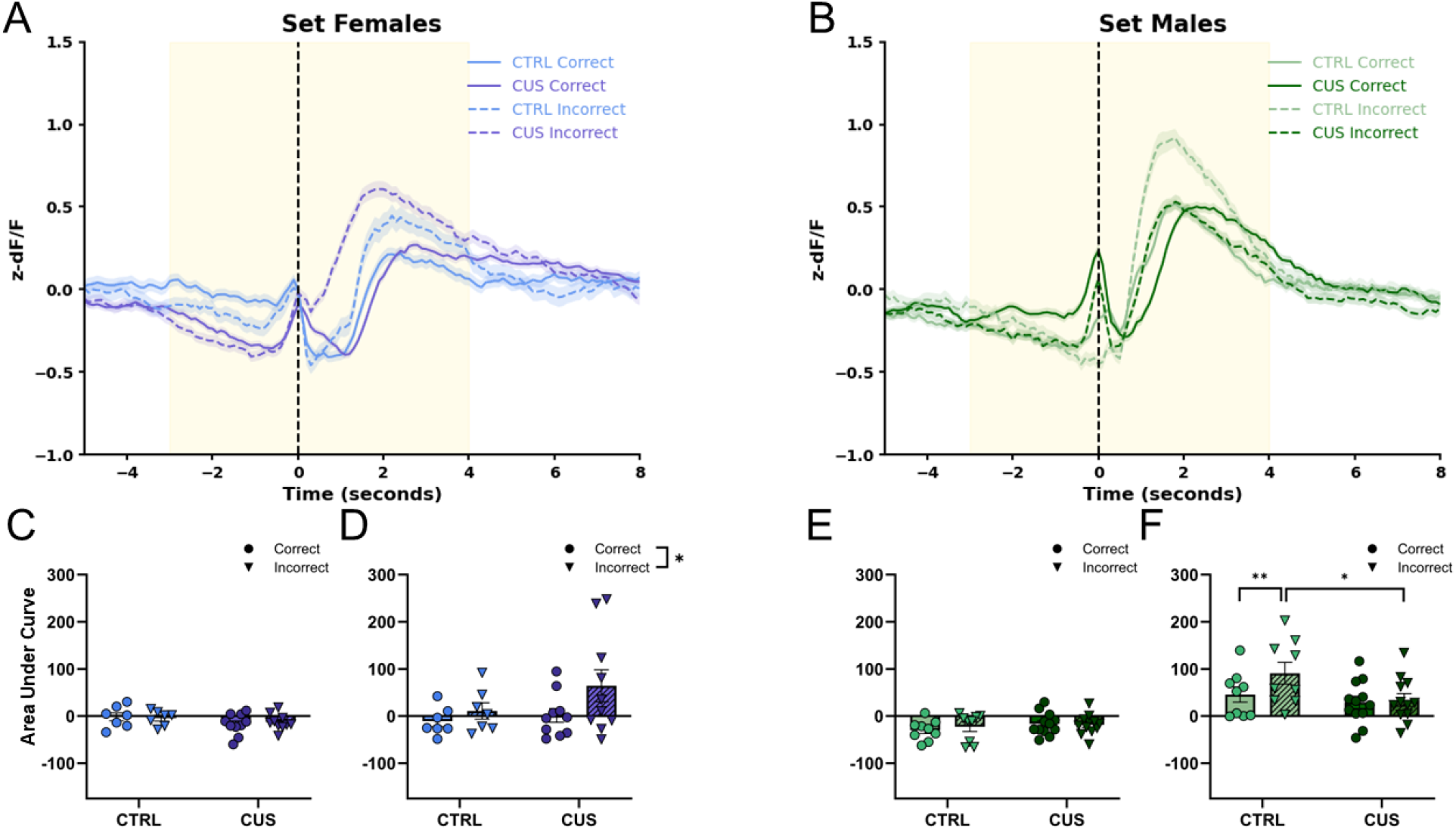
LHb Ca^2+^ dynamics during visual cue discrimination. Averaged group traces of visual cue discrimination-correct trials for **(A)** females and **(B)** males. Yellow shading indicates the data frames from which area under the curve (AUC) was calculated. AUC **(C)** pre- and **(D)** post-response for correct and incorrect trials during visual cue discrimination for females. AUC **(E)** pre- and **(F)** post-response for correct and incorrect trials during visual cue discrimination for males. Statistics are 3-way ANOVAs with sex, stress, and response type as factors. Main effects of response type are indicated in legend in the top right of each AUC graph. Main effects of sex are denoted in the graph area. Main effects of stress are common and not indicated in the graph. *p<0.05, **p<0.01,***p<0.001, ***<0.001)

